# Early-life acquisition of Lactobacillaceae protects against experimental cholera

**DOI:** 10.64898/2026.07.08.737341

**Authors:** Claire M. L. Chapman, Sophia Di Stefano, Andrew Kapinos Silva, Fabian Rivera-Chávez

## Abstract

Cholera causes severe diarrheal illness in young children, but the mechanisms underlying age-dependent susceptibility remain unclear. Experimental cholera in neonatal mice recapitulates age-dependent susceptibility: suckling mice are susceptible to *Vibrio cholerae* colonization and cholera toxin (CT)-dependent disease, whereas adult mice are not readily colonized and do not develop cholera-like disease. Here, we define a developmental window in which susceptibility declines sharply over the first two postnatal weeks. Maternal antibiotic exposure disrupted vertical transmission of maternal microbiota to offspring and altered distal small intestinal microbiota assembly, extending the window of susceptibility to CT-dependent *V. cholerae* colonization and disease. Pups born to antibiotic-treated dams exhibited reduced Lactobacillaceae and increased Enterobacteriaceae, and reintroduction of an endogenous *Lactobacillus* isolate restored offspring lactobacilli levels and reestablished resistance to experimental cholera at two weeks of age. Consistent with a direct protective role, increasing lactobacilli in susceptible neonatal mice reduced experimental cholera burden, and cultures of the endogenous *Lactobacillus* isolate as well as spent culture media acidified the *in vitro* growth environment and rapidly eliminated recoverable *V. cholerae*. Together, these findings identify vertical transmission of maternal microbiota to offspring and lactobacilli-associated antagonism as determinants of early-life resistance to cholera.

## INTRODUCTION

Young children are particularly susceptible to enteric infections that cause diarrheal disease, and cholera remains a major cause of severe diarrheal illness in children and adults worldwide^1,2^. Cholera is caused by the human pathogen *Vibrio cholerae* and is characterized by profuse secretory diarrhea that can be fatal within hours if left untreated. *V. cholerae* colonizes the small intestine, where it produces cholera toxin (CT), a potent enterotoxin that drives massive fluid loss. Since cholera is a human disease, conventional adult mice are not readily colonized by *V. cholerae* and do not develop cholera-like diarrheal disease. By contrast, experimental cholera can be modeled in suckling mice, which are highly susceptible to CT-dependent *V. cholerae* colonization and diarrheal-like disease^3,4^. This developmental transition provides an experimental system to investigate the mechanisms that define the age-dependent window of susceptibility and may be relevant to the heightened vulnerability of young children to diarrheal illness.

Early life is a critical developmental window during which neonates acquire intestinal microbiota from maternal and environmental sources that rapidly remodel the gut environment. During this period, immune system maturation and microbiota assembly are tightly linked such that perturbations in one can alter the development of the other^5,6^. The gut microbiota shapes mucosal immune development and function in the small intestine, influencing host resistance to enteric pathogen colonization and disease^7,8,9,6,10,11^.

Neonatal mammals acquire a significant portion of their intestinal microbiota through vertical transmission of maternal microbiota to offspring during birth and early life. Disruption of this process could alter postnatal community assembly and delay the acquisition of microbes that protect against enteric infection^12,13^. This question is particularly relevant because antibiotics are commonly administered during pregnancy^14^, and perinatal antibiotic exposure can disrupt vertical transmission of maternal microbiota to offspring and alter postnatal microbiota development^15,16,17^. Such perturbations may delay microbiota maturation and prevent acquisition of protective commensals that contribute to colonization resistance and protection from enteric disease. Here, we tested whether disruption of vertical transmission of maternal microbiota to offspring prolongs early-life susceptibility to experimental cholera.

## RESULTS

### Maternal antibiotic exposure delays age-dependent resistance to experimental cholera

To define the window over which susceptibility is lost in mice, we quantified age-dependent changes in *V. cholerae* colonization and CT-driven fluid accumulation in the distal small intestine, the primary site of *V. cholerae* colonization and disease. While it is well appreciated that suckling mice are highly susceptible to both *V. cholerae* colonization and CT-dependent diarrheal-like disease, whereas adult mice are resistant^3^, the age-dependent kinetics of susceptibility during postnatal development remain poorly defined. To define the developmental window in which mice lose susceptibility to experimental cholera, we orally infected C57BL/6J mice ranging from 5 days old through adulthood with wild-type *V. cholerae* C6706 or with the isogenic Δ*ctxAB* mutant (referred to hereafter as Δ*ctx*) lacking both the A and B subunits of CT, and quantified bacterial colonization in the distal small intestine and fluid accumulation in the gut. Wild-type *V. cholerae* colonized the distal small intestine at high levels in 5-day and 7-day-old mice, declined sharply in 14-day-old pups, and reached near the limit of detection in 21-day-old mice (Fig. 1A). Colonization by Δ*ctx* was significantly reduced relative to wild-type *V. cholerae* in susceptible 5-day-old mice, consistent with previous observations that the catalytic activity of CT promotes the pathogen’s growth in the small intestine of susceptible suckling mice^18^ (Fig. 1A). In parallel, wild-type *V. cholerae* infection induced diarrheal-like disease in 5-day- and 7-day-old mice, as indicated by significantly increased fluid accumulation relative to mock-infected controls, while this response progressively diminished by postnatal day 14 (Fig. 1B). In contrast, mice infected with the *V. cholerae* Δ*ctx* mutant did not develop diarrheal-like signs at any age, consistent with CT dependence in this model (Fig. 1B). In contrast, mice infected with the *V. cholerae* Δ*ctx* mutant did not devfigelop diarrheal-like signs at any age, consistent with CT dependence in this model (Fig. 1B). We observed a similar age-dependent decline in colonization and CT-dependent diarrheal disease in an outbred CD-1 background, supporting the generality of this developmental transition across mouse strains (Fig. S1A–B). Together, these data indicate that CT contributes to both colonization and disease during the first postnatal week, while CT-dependent disease and both CT-dependent and CT-independent colonization decline sharply by postnatal day 14.

**Figure 1.**
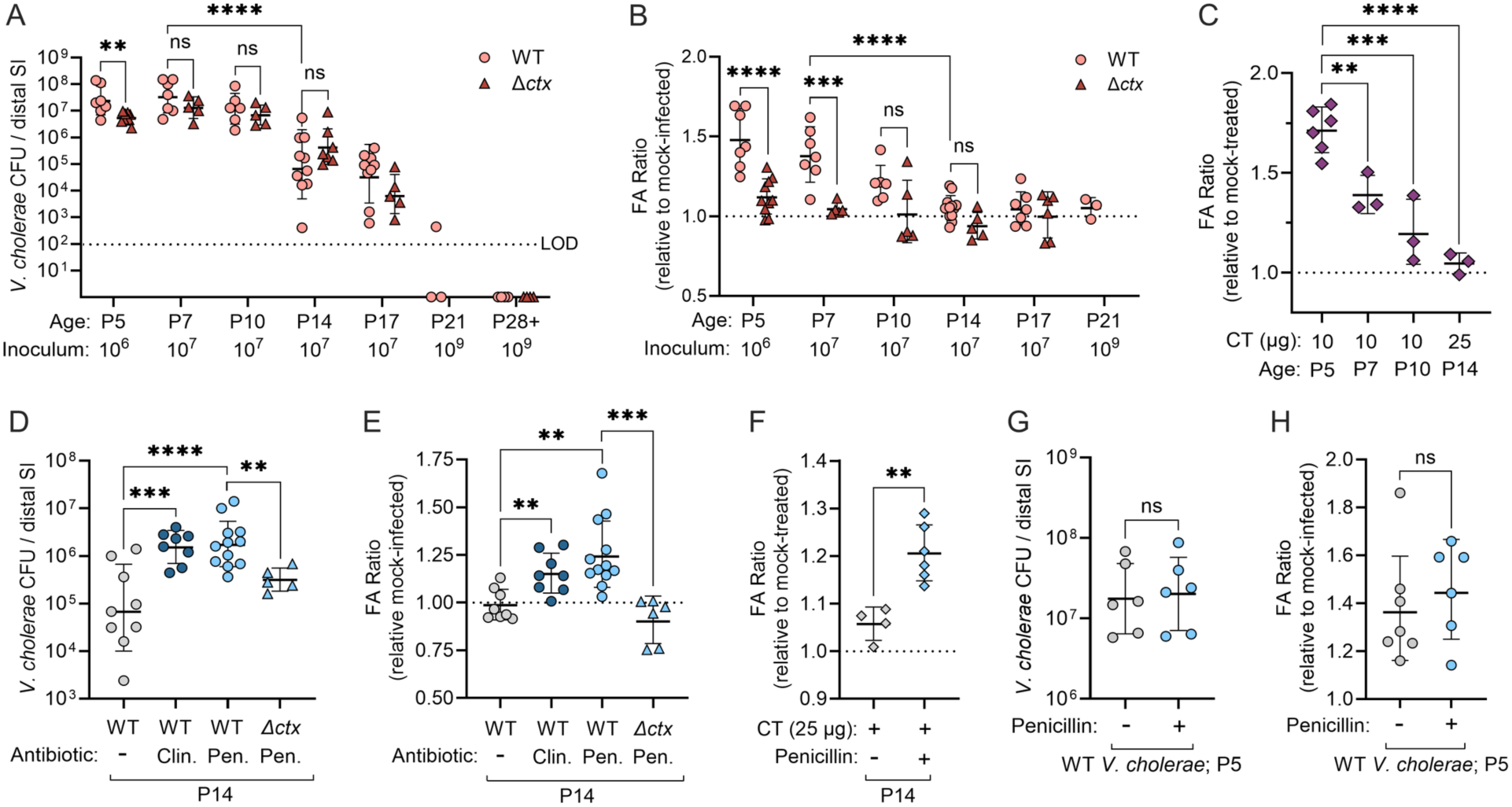
Maternal antibiotic exposure delays age-dependent resistance to experimental cholera. (A-B) C57BL/6J mice ranging from P5 to adulthood (P28+) were mock-infected with LB broth or infected with wild-type (WT) or CT-deficient *ΔctxAB V. cholerae mutant* (labeled *Δctx*). P5 mice received 10^6^ colony-forming units (CFU), P21 and P28+ mice received 10^9^ CFU, and all other groups received 10^7^ CFU. (A) Distal small intestinal *V. cholerae* CFU. The limit of detection (LOD) was 100 CFU, represented by the dotted line. (B) Fluid accumulation ratio relative to mock-infected controls. (C) C57BL/6J mice ranging from P5 to P14 were mock-treated with LB broth or orally treated with purified cholera toxin (CT). P14 mice received 25 μg CT and all other age groups received 10 μg CT. Fluid accumulation ratio is shown relative to mock-treated controls. (D–H) Breeding pairs received clindamycin (200 mg/L), penicillin (200 mg/L), or no antibiotics in drinking water throughout pregnancy and nursing. (D-E) P14 offspring were mock-infected with LB broth or infected with 10^7^ CFU WT or *ΔctxAB V. cholerae*. (D) Distal small intestinal *V. cholerae* CFU. (E) Fluid accumulation ratio relative to mock-infected controls. (G-H) P5 pups born to untreated or penicillin-treated breeding pairs were mock-infected with LB broth or infected with 10^6^ CFU WT *V. cholerae*. (G) Distal small intestinal *V. cholerae* CFU. (H) Fluid accumulation ratio relative to mock-infected controls. (F) Fluid accumulation ratio in P14 pups born to untreated or penicillin-treated breeding pairs following oral treatment with 25 μg purified CT relative to mock-treated controls. Each point represents an individual mouse. Bars indicate mean ± SEM. The dotted line in (A) indicates the limit of detection (LOD). The dotted lines in (B, E) indicate mock-infected controls, and the dotted lines in (C, F) indicate mock-treated controls. Statistical significance was determined by two-way ANOVA with Tukey’s multiple comparisons test in (A-B), ordinary one-way ANOVA with Dunnett’s multiple comparisons test in (C, E), ordinary one-way ANOVA with Šídák’s multiple comparisons test in (D), and unpaired t tests in (F-H) and where indicated. ns, not significant; *p < 0.05; **p < 0.01; ***p < 0.001; ****p < 0.0001.

We next asked whether the age-dependent decline in diarrheal-like disease reflects reduced expression of CT genes (*ctxAB*) by the pathogen or reduced host susceptibility to CT itself. To assess whether *V. cholerae* downregulates toxin expression in older animals, we measured *ctxAB* expression by qRT-PCR throughout the intestinal tract of wild-type *V. cholerae*-infected 14-day-old mice relative to 5-day-old mice. *ctxAB* expression was not reduced in 14-day-old mice compared to 5-day-old mice in any intestinal region tested (Fig. S1D). We then examined host responsiveness to the toxin by directly administering purified CT. Oral administration of purified CT induced robust fluid accumulation in 5-day-old mice, but this response waned with age and was no longer significantly increased in 14-day-old pups relative to mock-infected controls, despite 14-day-old mice receiving a higher CT dose than younger animals to account for increased body weight (Fig. 1C and Fig. S1C). Together, these results support a model in which developmental changes in the host limit CT-driven fluid accumulation by two weeks of age.

During early life, vertical transmission of maternal microbiota to offspring contributes to postnatal intestinal microbiota assembly and maturation^5,6^, raising the possibility that age-dependent resistance to *V. cholerae* colonization and CT-dependent fluid accumulation is linked to postnatal microbiota assembly. To examine this, we tested whether disrupting vertical transmission of maternal microbiota to offspring would prolong the window of susceptibility in mice. To this end, clindamycin or penicillin were administered in the drinking water of breeding pairs throughout pregnancy and nursing. 14-day-old pups born to antibiotic-treated breeding pairs were infected with wild-type *V. cholerae*. Maternal antibiotic exposure significantly increased colonization of wild-type *V. cholerae* in the distal small intestine of 14-day-old offspring relative to pups born to untreated dams (Fig. 1D). Additionally, 14-day-old pups born to antibiotic-treated dams and infected with wild-type *V. cholerae* also exhibited significantly increased fluid accumulation compared to pups born to untreated dams (Fig. 1E). We focused subsequent experiments on penicillin because it produced a robust increase in both *V. cholerae* colonization and fluid accumulation (Fig. 1D–E) and because penicillin and similar beta-lactam antibiotics are widely used during pregnancy^19,20^. Since antibiotic exposure itself can affect intestinal physiology, we tested whether the increased fluid accumulation in pups born to penicillin-treated dams remained dependent on the catalytic action of CT. In contrast to wild-type infection, 14-day-old pups born to penicillin-treated dams and infected with the *V. cholerae* Δ*ctx* mutant did not develop diarrheal-like signs, as measured by fluid accumulation, demonstrating that the disease phenotype remained CT dependent (Fig. 1E). Consistent with increased host responsiveness to CT, 14-day-old pups born to penicillin-treated dams exhibited significantly increased fluid accumulation following oral treatment with purified CT compared with pups born to untreated dams (Fig. 1F).

Previous work has shown that the catalytic activity of CT promotes *V. cholerae* growth in the small intestine of susceptible neonatal animals^18,21,22,23^. Since maternal penicillin treatment prolonged susceptibility to CT-dependent disease in 14-day-old pups, we next asked whether it also restored the CT-dependent growth advantage of *V. cholerae*. In 14-day-old pups born to penicillin-treated dams, wild-type *V. cholerae* was recovered at significantly higher levels than the Δ*ctx* mutant (Fig. 1D), whereas no significant difference between the two strains was observed in 14-day-old pups born to untreated dams (Fig. 1A), further supporting the conclusion that disrupted vertical transmission of maternal microbiota to offspring prolongs susceptibility to experimental cholera into the second postnatal week.

Antibiotic-mediated disruption of the gut microbiota can reduce colonization resistance against enteric pathogens^24^. We therefore asked whether maternal penicillin further reduced colonization resistance during the already susceptible neonatal window. To test this, we infected 5-day-old pups born to untreated or penicillin-treated breeding pairs with wild-type *V. cholerae*. Maternal penicillin did not significantly alter *V. cholerae* colonization or fluid accumulation in susceptible 5-day-old pups (Fig. 1G–H). Together, these findings demonstrate that maternal antibiotic exposure prolongs the early-life window of susceptibility to *V. cholerae* colonization and CT-dependent disease, and that maternal penicillin treatment increases host responsiveness to the intestinal action of CT.

### Maternal penicillin disrupts neonatal distal small intestinal microbiota assembly and reconstitution of endogenous *Lactobacillus* restores resistance to experimental cholera

Maternal penicillin prolonged susceptibility to experimental cholera, suggesting that altered development of the intestinal microbiota plays a role in the extended susceptibility window. To examine how maternal penicillin treatment alters the microbiota in the distal small intestine of neonatal mice, where *V. cholerae* colonizes and produces CT, we performed 16S rRNA gene sequencing of distal small intestinal contents from 5- and 14-day-old pups born to penicillin-treated or untreated dams, together with fecal samples from the corresponding adult dams. In pups born to untreated dams, the distal small intestinal microbiota underwent a marked developmental transition between 5 and 14 days of age, characterized at the family level by increased relative abundance of Lactobacillaceae and a decline in Enterobacteriaceae (Fig. 2A). This developmental enrichment of *Lactobacillus* is consistent with prior observations of normal postnatal maturation of the murine small intestinal microbiota, with *Lactobacillus* enriched by 2–3 weeks of age^25^. CLR-normalized abundance analyses confirmed that maternal penicillin markedly reduced Lactobacillaceae in both 5-day- and 14-day-old pups (Fig. 2B–C) and in fecal samples from corresponding dams (Fig. 2D). In pups born to penicillin-treated dams, Enterobacteriaceae expanded and dominated the community at both ages (Fig. 2A and Fig. S2E), consistent with previous observations that antibiotic perturbation can disrupt intestinal microbial community structure and favor expansion of Enterobacteriaceae^26^. Although individual mice varied, particularly among penicillin-treated pups, order- and class-level profiles, individual family-level profiles, and relative-abundance analyses showed the same overall age- and penicillin-associated shift away from Lactobacillaceae and toward Enterobacteriaceae (Fig. S2A–E). Together, these data indicate that maternal penicillin disrupts developmental assembly of the neonatal small intestinal microbiota, with a pronounced loss of Lactobacillaceae-associated taxa.

**Figure 2.**
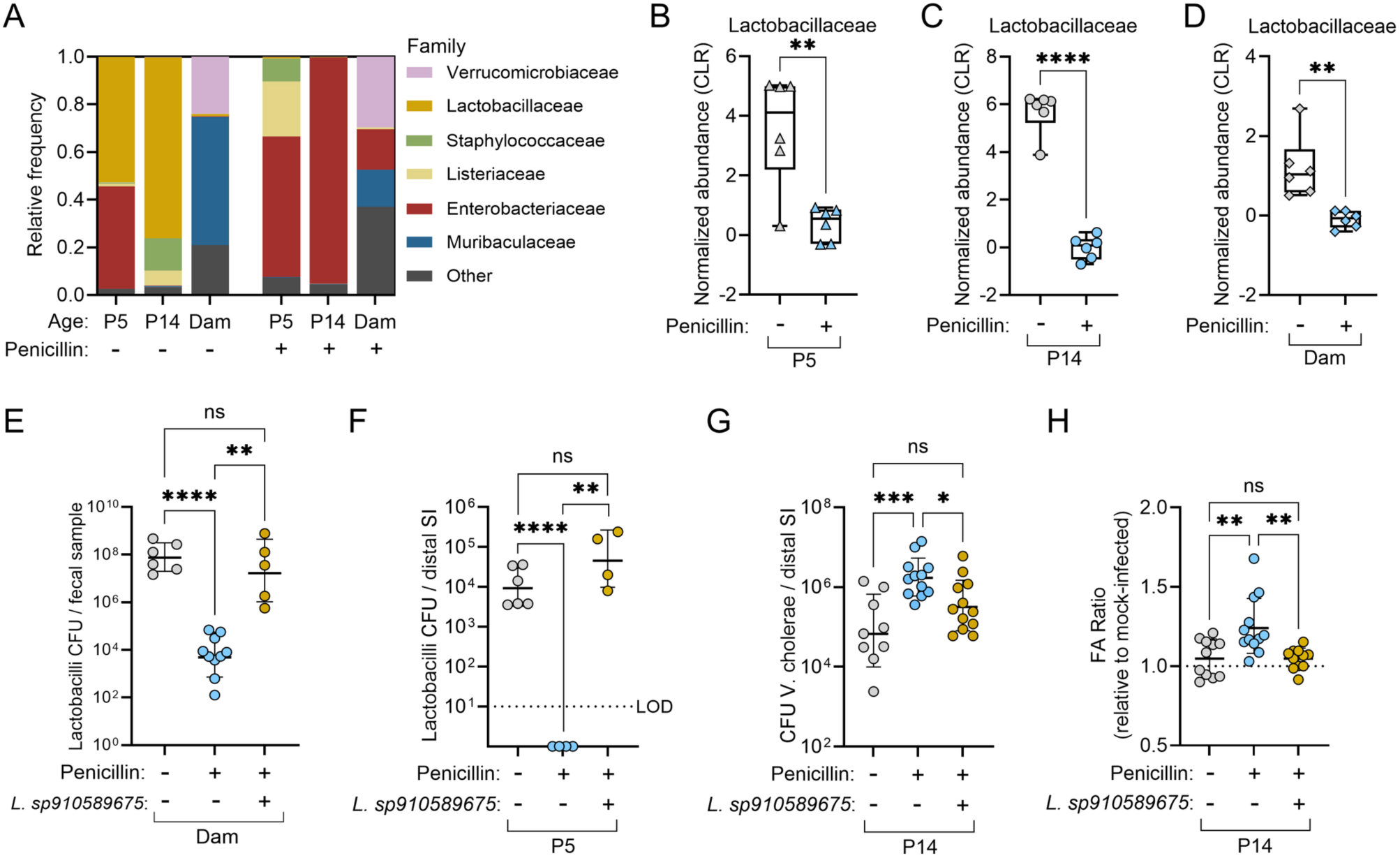
Maternal penicillin disrupts neonatal distal small intestinal microbiota assembly and reconstitution of endogenous *Lactobacillus* restores resistance to experimental cholera. (A–D) Distal small intestinal bacterial communities from P5 and P14 pups born to untreated or penicillin-treated breeding pairs, together with fecal communities from the corresponding adult dams, were analyzed by 16S rRNA gene sequencing. (A) Family-level relative abundance of distal small intestinal bacterial communities. (B-D) CLR-normalized abundance of *Lactobacillaceae* in P5 pups, P14 pups, and dams, respectively. (E–H) Penicillin-treated breeding pairs received *Lactobacillus sp910589675*, an endogenous strain isolated from 21-day-old mice. (E) Recoverable *lactobacilli* from fecal samples of untreated dams, penicillin-treated dams, or penicillin-treated dams receiving *Lactobacillus sp910589675*. (F) Recoverable *lactobacilli* CFU in the distal small intestine of P5 pups born to untreated dams, penicillin-treated dams, or penicillin-treated dams receiving *Lactobacillus sp910589675*. (G) Distal small intestinal *V. cholerae* CFU and (H) fluid accumulation ratio in P14 pups from the indicated maternal treatment groups following infection with 10⁷ CFU WT *V. cholerae*. Each point in (B–H) represents an individual biological sample or mouse, as appropriate. Bars indicate mean ± SEM. Statistical significance in (B-D) was determined by Welch’s t test. Statistical significance in (E-F) was determined by Brown-Forsythe and Welch ANOVA with Dunnett’s T3 multiple comparisons test. Statistical significance in (G–H) was determined by ordinary one-way ANOVA with Šídák’s multiple comparisons test. ns, not significant; *p < 0.05; **p < 0.01; ***p < 0.001; ****p < 0.0001.

The pronounced depletion of Lactobacillaceae from the distal small intestinal microbiota of neonatal pups following maternal penicillin treatment identified this family as a candidate contributor to age-dependent resistance to experimental cholera. Notably, *Lactobacillus* is a dominant genus in the murine small intestine, and several species within this genus are well-characterized probiotic strains with protective activity against enteric pathogens^27,28^. We therefore sought to examine whether restoring endogenous *Lactobacillus* species could reverse the antibiotic-extended susceptibility phenotype. To this end, we isolated endogenous *Lactobacillus* from the small intestinal contents of 21-day-old mice and identified *Lactobacillus sp910589675*. We then orally administered this isolate to penicillin-treated breeding pairs during pregnancy and nursing to test whether restoration of a native *Lactobacillus* strain could reverse antibiotic-extended susceptibility to experimental cholera in their offspring.

Direct bacterial enumeration on selective MRS media confirmed that maternal penicillin depleted recoverable *Lactobacillus* from fecal samples of dams. Oral administration of *Lactobacillus sp910589675* to penicillin-treated breeding pairs significantly restored fecal *Lactobacillus* levels to those observed in untreated dams (Fig. 2E). Administration of *Lactobacillus sp910589675* to breeding pairs also restored recoverable *Lactobacillus* in 5-day-old pups born to penicillin-treated dams to levels comparable to those observed in pups born to untreated dams (Fig. 2F). To test whether reestablishing endogenous *Lactobacillus* levels was sufficient to restore resistance to experimental cholera, we infected 14-day-old pups born to untreated dams, penicillin-treated dams, or penicillin-treated dams that received the *Lactobacillus* add-back with wild-type *V. cholerae*. Reestablishing endogenous *Lactobacillus* levels reduced *V. cholerae* colonization relative to penicillin-treated controls (Fig. 2G) and reduced fluid accumulation in *V. cholerae*–infected pups to levels comparable to those observed in 14-day-old pups born to untreated dams (Fig. 2H). Collectively, these findings demonstrate that disruption of early-life *Lactobacillus* acquisition contributes to the antibiotic-extended susceptibility window and that restoring vertical transmission of endogenous *Lactobacillus* is sufficient to reduce both *V. cholerae* colonization and CT-dependent disease.

### Acute elevation of lactobacilli reduces experimental cholera in susceptible suckling mice

Having found that maternal administration of an endogenous *Lactobacillus* strain restored resistance to experimental cholera in antibiotic-exposed 14-day-old pups, we next asked whether increasing recoverable lactobacilli in the small intestine during the otherwise susceptible neonatal window could also provide protection. Since 5-day-old suckling mice naturally harbor lower levels of lactobacilli than 14-day-old mice (Fig. 2), we orally gavaged 4-day-old mice with *Ligilactobacillus murinus*, MRS broth alone, or LB broth alone as a mock treatment and directly enumerated recoverable lactobacilli 24 hours later. Oral administration of *L. murinus* produced a trend toward increased recoverable lactobacilli in 5-day-old mice that did not reach significance (Fig. 3A). Unexpectedly, treatment with MRS broth alone significantly increased recoverable lactobacilli, suggesting that MRS may promote expansion of endogenous lactobacilli in the small intestine (Fig. 3A). We next asked whether these pretreatments reduced susceptibility to experimental cholera. Mice were infected at 5 days of age with wild-type *V. cholerae*. Pretreatment with *either L. murinus* or MRS broth significantly reduced *V. cholerae* colonization in the distal small intestine compared with untreated controls (Fig. 3B) and significantly reduced fluid accumulation after infection (Fig. 3C). These findings indicate that pretreatment with either *L. murinus* or MRS broth reduces susceptibility to experimental cholera in otherwise susceptible suckling mice.

**Figure 3.**
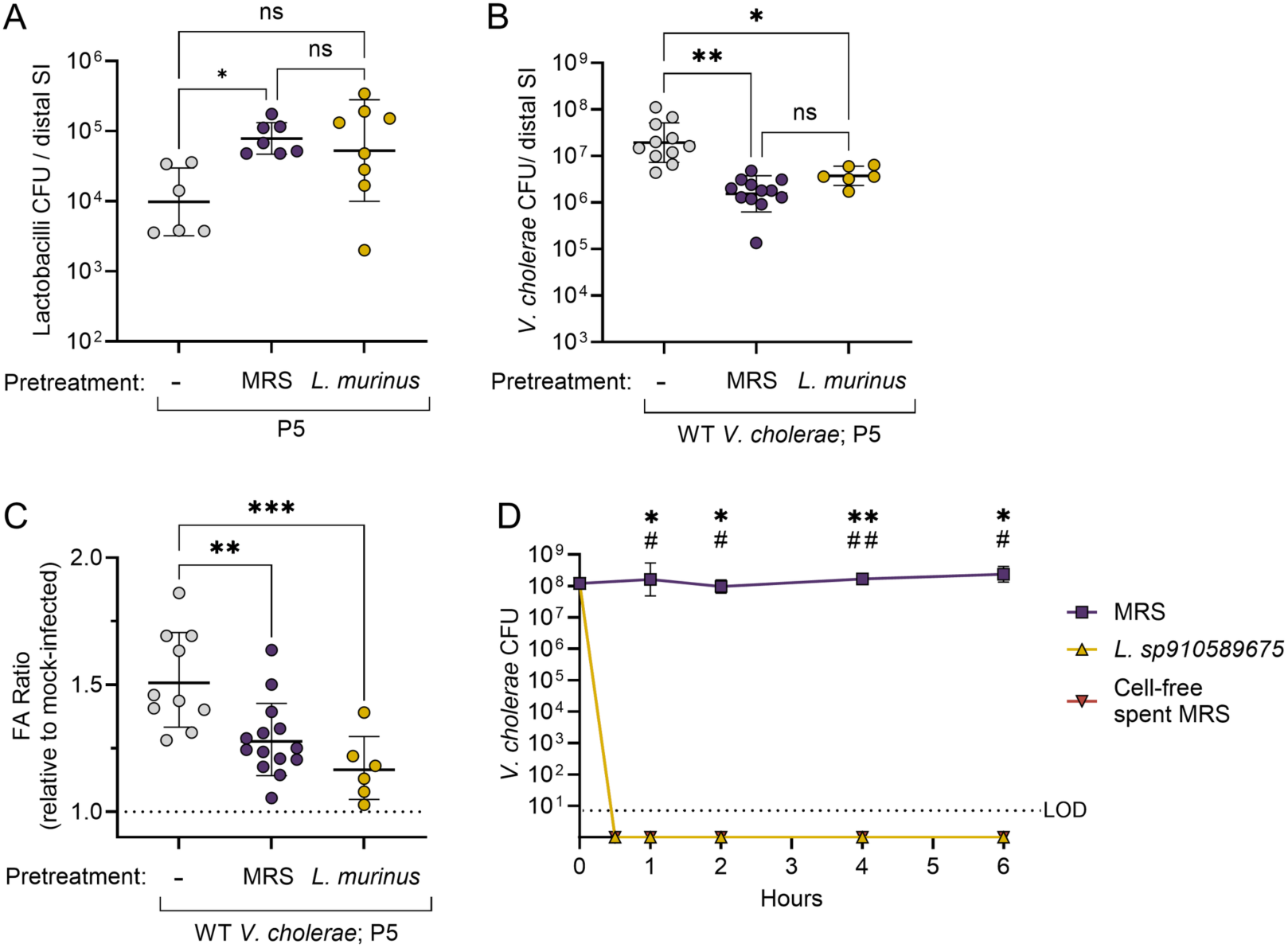
Acute elevation of lactobacilli reduces experimental cholera in susceptible suckling mice. (A-C) P4 C57BL/6J mice were pretreated with *L. murinus* or MRS broth and analyzed 24 hours later at P5 (A) or mock-infected with LB broth or infected with 10^6^ CFU WT *V. cholerae* (B-C). (A) Recoverable *Lactobacillus* CFU in the distal small intestine following 24 hours of the indicated pretreatments. (B) Distal small intestinal *V. cholerae* CFU in P5 mice following the indicated pretreatments and infection with WT *V. cholerae*. (C) Fluid accumulation ratio relative to mock-infected mice receiving the same pretreatment. (D) Recovery of *V. cholerae* following incubation in fresh MRS broth, an MRS-grown *Lactobacillus sp910589675* culture, or cell-free spent MRS medium for the indicated times. No recoverable *V. cholerae* colonies were detected after exposure to the *Lactobacillus* culture or cell-free spent MRS medium (these values are plotted below the limit of detection (LOD) for visualization). Each point in (A–C) represents an individual mouse. Bars indicate mean ± SEM. Dotted line in (C) represents mock-infected controls. Statistical significance in (A) was determined by Brown-Forsythe and Welch ANOVA with Dunnett’s T3 multiple comparisons test. Statistical significance in (B) was determined by ordinary one-way ANOVA with Šídák’s multiple comparisons test. Statistical significance in (C) was determined by ordinary one-way ANOVA with Tukey’s multiple comparisons test. Statistical significance in (D) was determined by Kruskal-Wallis test with Dunn’s multiple comparisons test at each indicated time point. Asterisks (**\***) denote comparisons between fresh MRS broth and the *L. sp910589675* culture, and pound signs (#) denote comparisons between fresh MRS broth and the cell-free spent MRS medium. ns, not significant; *p < 0.05; **p < 0.01; ***p < 0.001.

To determine whether products generated during growth of the endogenous *Lactobacillus* isolate, including lactic acid, could directly antagonize *V. cholerae*, we incubated V. cholerae in fresh MRS broth, an MRS-grown *Lactobacillus sp910589675* culture, or cell-free spent MRS medium. Fresh MRS had a pH of approximately 6, whereas both the *Lactobacillus* culture and spent medium were acidified to approximately pH 4. While *V. cholerae* remained recoverable in fresh MRS broth, exposure to the *Lactobacillus* culture or spent medium significantly reduced recoverable *V. cholerae* to below the limit of detection within 30 minutes (Fig. 3D). These findings suggest that acidification during growth of the endogenous *Lactobacillus* isolate, potentially through production of lactic acid or other organic acids, contributes to the rapid loss of *V. cholerae*. Together, these results demonstrate that pretreatment with *L. murinus* or MRS broth reduces experimental cholera in fully susceptible suckling mice and that products generated during growth of the endogenous *Lactobacillus* isolate directly inhibit *V. cholerae*.

These findings identify vertical transmission of the maternal microbiota to offspring as a determinant of age-dependent resistance to experimental cholera and support lactobacilli as protective early-life commensals that limit *V. cholerae* through both colonization resistance and rapid direct antagonism.

## DISCUSSION

Previous studies have linked gut microbiota composition to susceptibility to cholera in humans and to protection from *V. cholerae* colonization in mouse models. In household contacts of patients with cholera, younger age was associated with increased susceptibility to infection, and fecal microbiota composition was also associated with both age and subsequent susceptibility^29^. These observations raised the possibility that developmental changes in the intestinal microbiota contribute to age-dependent cholera susceptibility but did not establish a causal mechanism. In a separate study, bacterial taxa that increased during recovery from cholera in adults were incorporated into a defined human gut community in gnotobiotic mice, in which *Ruminococcus obeum* restricted *V. cholerae* colonization^30^. However, *Ruminococcus obeum* is primarily a colonic bacterium isolated from stool samples of cholera patients. Thus, whether members of the microbiota from the small intestine, where *V. cholerae* colonizes and produces CT, participate in the resistance to cholera remained an important unanswered question. Furthermore, whether vertical transmission of maternal microbiota to offspring and early-life microbiota assembly contribute to the emergence of resistance to experimental cholera as mice age has not been examined.

Our findings show that mice acquire resistance to experimental cholera before weaning, by two weeks of age. Maternal antibiotic treatment delayed this developmental transition by disrupting assembly of the small intestinal microbiota. Although neonatal small intestinal communities are low-diversity at baseline, maternal penicillin produced a pronounced compositional shift characterized by reduced Lactobacillaceae and increased Enterobacteriaceae in the distal small intestine of neonatal offspring. Antibiotic-associated Enterobacteriaceae blooms have been linked to disruption of colonization resistance and inflammation-dependent metabolic changes, including nitrate respiration by commensal *E. coli* after streptomycin treatment^31^. In our model, however, increased Enterobacteriaceae occurred alongside increased *V. cholerae* colonization and CT-dependent disease. This is consistent with prior work showing that *V. cholerae* can antagonize commensal bacteria through its type VI secretion system (T6SS), facilitating colonization in the presence of competing Enterobacteriaceae^32^. Although these findings do not distinguish whether the Enterobacteriaceae expansion directly contributes to enhanced *V. cholerae* colonization, this possibility appears less likely given that *E. coli* limits colonization of T6SS-deficient *V. cholerae* but not strains that harbor an intact T6SS^32^. Thus, the increase in Enterobacteriaceae likely reflects antibiotic-associated dysbiosis, although its specific contribution to infection in this model remains to be determined.

Previous studies have shown that *Lactobacillus* becomes a dominant component of the murine small intestinal microbiota during postnatal maturation^25^. The pronounced depletion of Lactobacillaceae in offspring of penicillin-treated breeding pairs in this study implicated *Lactobacillus* as a candidate mediator of age-dependent resistance to experimental cholera. While penicillin-treated dams retained a mixed fecal community, their two-week-old pups developed an Enterobacteriaceae-dominant distal small intestinal microbiota with marked depletion of Lactobacillaceae. Since pups were not directly administered antibiotics, these findings indicate that maternal penicillin treatment leads to a persistent perturbation of the small intestinal microbiota in offspring. Although administration of the lactic acid-producing bacterium *Lactococcus lactis* can reduce *V. cholerae* burden in infant mice^33^, the role of endogenous *Lactobacillus* in the emergence of resistance to experimental cholera in neonatal mice, particularly CT-dependent diarrheal-like disease, had not previously been defined. Given its established roles in neonatal intestinal maturation, *Lactobacillus* was a strong candidate to shape age-dependent resistance.

Restoring endogenous *Lactobacillus* in penicillin-treated breeding pairs reestablished *Lactobacillus* levels in their offspring and restored resistance to both *V. cholerae* colonization and CT-dependent fluid accumulation in two-week-old pups. These findings establish that impaired vertical transmission of *Lactobacillus* from mothers to offspring contributes functionally to the antibiotic-extended susceptibility window.

The protective activity of lactobacilli was also evident after acute administration in otherwise susceptible suckling mice. Similar to prior findings with lactic acid-producing bacterium *Lactococcus lactis*^33^, both *L. murinus* and MRS broth reduced *V. cholerae* colonization and fluid accumulation when administered before infection. *In vitro*, growth of *Lactobacillus* acidified MRS medium from approximately pH 6.4 to pH 4, and both the MRS-grown *Lactobacillus* culture and cell-free spent medium eliminated recoverable *V. cholerae* within 30 minutes, whereas MRS broth did not. These findings support a model in which products generated during *Lactobacillus* growth directly antagonize *V. cholerae*. Although the active factor remains to be defined, the accompanying acidification suggests that *Lactobacillus*-derived organic acids may contribute to the rapid loss of recoverable bacteria.

Direct antagonism may explain the reduction in *V. cholerae* colonization but is unlikely to fully account for the restored resistance to CT-dependent fluid accumulation. We therefore propose that lactobacilli protect through at least two separable mechanisms: direct inhibition of *V. cholerae* growth and modulation of host pathways that regulate susceptibility to CT-driven fluid secretion. One possible host mechanism may involve *Lactobacillus*-dependent support of epithelial homeostasis. Recent studies showed that *Lactobacillus*-derived phenyllactic acid can activate epithelial PPARγ signaling^34^, a pathway linked to preservation of epithelial metabolic homeostasis and restriction of dysbiotic Enterobacteriaceae expansion^35^. Depletion of Lactobacillaceae after maternal penicillin treatment could therefore reduce microbiota-derived PPARγ agonists and weaken epithelial programs that normally limit Proteobacteria expansion and support mucosal resilience. Another possible host mechanism may involve microbiota-dependent maturation of IL-22-associated epithelial programs and ion transport pathways, including SLC26A3, which have been linked to protection from dehydration and fluid loss during enteric infection^36,37,38^. Determining whether endogenous *Lactobacillus* promotes these epithelial responses will be an important direction for future studies.

More broadly, these findings identify vertical transmission of maternal microbiota to offspring as a determinant of neonatal small intestinal microbiota assembly and the early-life emergence of resistance to experimental cholera. They further identify lactobacilli as a functional contributor to this transition, linking early-life microbiota assembly to *V. cholerae* colonization, CT-dependent disease, and host susceptibility to the intestinal action of CT. In breastfed human infants, the fecal microbiota is commonly dominated by *Bifidobacterium*, which shares relevant early-life functions with *Lactobacillus*^39,40^. Thus, this developmental host-microbe mechanism may be relevant to the heightened susceptibility of young children to cholera and other diarrheal diseases.

## ACKNOWLEDGMENTS

We thank members of the Rivera-Chávez lab for insightful discussions and technical support. We thank Dr. Rob Knight and members of the Knight laboratory at UC San Diego for expert technical assistance with microbiota sequencing analysis. C.M.L.C. is funded by the National Science Foundation Graduate Research Fellowship Program. F.R.-C. is funded by UC San Diego, the Edward Mallinckrodt Jr. Foundation, the Hellman Family Foundation, and the UC San Diego/San Diego Digestive Diseases Research Center (DK120515).

## DECLARATION OF INTERESTS

The authors declare no competing interests.

## AUTHOR CONTRIBUTIONS

F.R.-C. and C.M.L.C., designed and conceived the study; C.M.L.C., S.D.S., and A.K.S. performed the experiments and data analysis. F.R.-C. and C.M.L.C. wrote the manuscript.

## METHODS

### Bacterial strains

Bacterial strains used in this study are listed in Table S1. Unless indicated otherwise, *V. cholerae* strains were routinely grown aerobically at 37°C in LB broth or on LB plates for 15 hours. *Ligilactobacillus murinus and Lactobacillus sp910589675* were routinely grown anaerobically at 37°C in de Man, Rogosa and Sharpe (MRS) broth (BD Difco # 288130) or on MRS agar (BD Difco #288210) plates using an AnaeroPack 7.0L Rectangular Jar (Thermo Scientific # 23-246-387) and 3 AnaeroPack-Anaero Anaerobic Gas Generators (Thermo Scientific # 23-246-376) for 24 hours. When indicated, antibiotics were added to the media at the following concentrations: 100 μg/ml streptomycin (*V. cholerae*).

### Animal Experiments

All animal experiments in this study were approved by the Institutional Animal Care and Use Committee at University of California, San Diego.

### Mouse Experiments

Suckling mice (P4-P21) were separated from their dams 1 hour prior to infection. Mice were randomly allocated to treatment groups and orally inoculated with 1 × 10^6^ CFU (C57BL/6J, P5), 1 × 10^7^ CFU (CD-1 P5-P17, C57BL/6J P7-P17), or 1 × 10^9^ CFU (C57BL/6J P21) *V. cholerae,* 50 μl sterile LB broth (mock infection), or 10 μg (C57BL/6J, P5-P10) or 25 μg (C57BL/6J, P14) of purified CT (List Biological Laboratories, Campbell, CA). CD-1 mice were obtained from Charles River Laboratories (Wilmington, MA, USA) and C57BL/6J mice were bred in-house.

Adult CD-1 and C57BL/6J mice were 4 weeks old at the time of infection. Adult mice were randomly allocated to treatment groups and orally inoculated with 1 × 10^9^ CFU *V. cholerae* or 100 μl sterile LB broth (mock infection).

Mice were placed inside a 30°C incubator. After 22 hours, mice were euthanized and the gastrointestinal tracts were aseptically removed. The proximal half (by length) of the small intestine was defined as the “proximal small intestine” (pSI) and the distal half of the small intestine was defined as the “distal small intestine” (dSI). To measure bacterial burden, entire pSI, dSI, and colon regions were homogenized in 1mL sterile PBS with scissors, vortexed for one minute. Serial dilutions were plated on LB agar supplemented with 100 μg/mL streptomycin to enumerate *V. cholerae* or on MRS agar to enumerate recoverable *lactobacilli*. Plates were incubated overnight under aerobic (LB) or anaerobic (MRS) conditions at 37°C. For DNA and RNA extraction pSI, dSI, and colon contents were squeezed out and both contents and tissue were snap frozen in liquid nitrogen and stored at -80°C until use.

The fluid accumulation (FA) ratio was calculated by weight of the entire gastrointestinal tract (gut) / (mouse body weight - gut weight). Animals that were euthanized before an experimental endpoint due to health concerns were excluded from the analysis.

### RNA Extraction and Quantitative RT-PCR (qRT-PCR)

To extract RNA from snap frozen tissue, tissue was resuspended in 1mL TRIzol reagent (Invitrogen # 15596018) in Green RINO RNA bead lysis kit (Next Advance, #GREENR1-RNA) and homogenized for 5 minutes (p5 tissue) or 10 minutes (p14-adult tissue) in Bullet Blender Storm Pro homogenizer, speed 10. Alternatively, to extract RNA from snap frozen intestinal contents, contents were resuspended in 1mL TRIzol reagent. 200 μl of chloroform was added to content/tissue and TRIzol mixture (and removed from bead tube if necessary), mixed, and incubated at room temperature for 2 minutes. Samples were then centrifuged at 15000xg for 15 minutes at 4°C and the aqueous phase was transferred to a new tube. 400 μl of 70% ethanol was added to samples then transferred to a RNeasy micro kit (QIAGEN, #74004) column and the isolation protocol was continued according to manufacturer’s instructions with the exception that samples were eluted in UltraPure Distilled water (Invitrogen, #10977-015) rather than the provided elution buffer. The concentration of the RNA was determined using Nanodrop One and samples were diluted to 10 ng/μl and stored at -80°C until used.

qRT-PCR was performed using SuperScript III Platinum One-Step qRT-PCR master mix (Invitrogen, #12574026) according to the manufacturer’s instructions. Each sample had two technical replicates and 2 μl of 10ng/μl RNA template in a 20μl reaction per target gene. A list of target gene primers can be found in Table S1. Samples were run on a CFX Opus 384 Real-time PCR thermocycler (BioRad) cycle info. No-template and no-reverse-transcriptase controls were run for each target gene on each plate. Gene expression was normalized to a housekeeping gene (*ActB* for mouse tissue, gyrB for *V. cholerae*) and control group using the 2^−ΔΔCt method^41^.

### Perinatal Antibiotic Administration

Pregnant dams were given clindamycin (200 mg/L), penicillin (200 mg/L), or no antibiotic in their drinking water for the duration of their first 4 litters. Dam fecal samples were collected at the time of each experiment and diluted and plated onto MRS agar to measure *Lactobacillus* abundance.

### Microbiota Sequencing and Analysis

Distal small intestinal contents from pups and fecal samples from dams were collected at the indicated experimental time points and stored at −80°C until processing. Microbiota profiling was performed by the UC San Diego Center for Microbiome Innovation/Knight laboratory using 16S rRNA gene amplicon sequencing as described^42^. Briefly, DNA was extracted from intestinal contents or fecal samples, and bacterial 16S rRNA gene amplicon libraries targeting the V4 region were generated using a high-throughput amplicon sequencing workflow based on the Earth Microbiome Project/Knight laboratory protocol. Amplicons were sequenced on an Illumina platform.

Demultiplexed sequence data were processed using QIIME 2^43^. Briefly, amplicons were demultiplexed and quality control was performed using default workflows, followed by trimming to 100 bp and taxonomy selection using a closed OTU model (Greengenes 13_8 97% identity OTUs)^44^. Microbial α-diversity and alpha-rarefaction curves were measured in QIIME 2 using Faith’s Phylogenetic Diversity^45^ and Shannon index^46^. For preliminary comparisons of all results, a sampling depth of approximately 1,000 reads per sample was selected; in subsequent analyses, the output from P5 animals was sampled at a depth of 211 reads per sample to account for reduced α-diversity and sampling depth in penicillin-treated neonates. β-diversity was calculated using Bray-Curtis dissimilarity^47^, followed by principal coordinate analysis (PCoA) using default QIIME 2 parameters. Relative abundance plots were generated from taxonomic summaries, and centered log-ratio (CLR)-normalized abundances were calculated using R version 4.5.3 and the packages “compositions” (v.2.0.9)^48^ and “biomformat” (v.1.38.3)^49^.

Statistical analyses were performed in GraphPad Prism 8.0 (GraphPad Software, La Jolla, CA). For multiple comparisons, Brown-Forsythe and Welch ANOVA tests were used. For single comparisons, Welch’s t-test was used. In both types of comparison, a p-value threshold of 0.05 was applied.

### Lactobacillus sp910589675 reconstitution

*Lactobacillus sp910589675* was recovered by plating small-intestinal contents from a 21-day-old female C57BL/6J mouse on MRS agar. Three colonies were isolated and identified by whole-genome sequencing as *Lactobacillus sp910589675*. Bacterial genome sequencing was performed by Plasmidsaurus using Oxford Nanopore Technology with custom analysis and annotation.

To reconstitute breeding pairs, *L. sp910589675* was grown anaerobically at 37°C in MRS broth for 24 hours. The culture was pelleted and resuspended in MRS broth to a concentration of 1×10^10^ CFU/mL. Mice in each breeding were orally gavaged with 1×10^9^ CFU *L. sp910589675* every other day from gestational day 7 through the experimental endpoint. Fecal samples from breeders were plated on MRS agar to quantify recoverable *Lactobacillus*. 5- or 14-day-old pups born to these breeding pairs were infected or mock-infected as described above.

### Acute Ligilactobacillus Addition

*Ligilactobacillus murinus* was grown anaerobically at 37°C in MRS broth. After 24 hours, the culture was pelleted and resuspended in LB broth to minimize carryover of MRS. P4 mice were orally gavaged with 1×10^9^ CFU *L. murinus* suspended in LB broth, MRS broth alone, or LB broth alone as a mock treatment and returned to the dam. 24 hours later, pups were separated from dams for 1 hour and either euthanized for *Lactobacillus* enumeration or mock-infected or infected with wild-type *V. cholerae* as described above.

### *In vitro* Growth Assay

*Lactobacillus sp910589675* was grown anaerobically in MRS broth for 24 h. Half of the culture was retained as an intact *Lactobacillus* culture, and the remaining half was pelleted and filtered through a 0.22-μm filter (Millipore #SLGPR33RB) to generate cell-free spent MRS medium. *V. cholerae* was incubated at 37°C in fresh MRS broth, an MRS-grown *Lactobacillus sp910589675* culture, or cell-free spent MRS medium for 0.5, 1, 2, 4, or 6 h. At inoculation and each indicated time point, cultures were plated on LB agar supplemented with 100 μg/mL streptomycin to enumerate recoverable V. cholerae. 100 μg/ml streptomycin to measure recoverable *V. cholerae*. The pH of each condition was measured qualitatively with pH strips (Defirm SC001).

### Statistical Analysis

Statistical analyses were performed using GraphPad Prism version 10. When comparing two groups, an unpaired t-test was performed for parametric data and a Mann-Whitney test for non-parametric data. When comparing one variable across three or more-groups, a one-way ANOVA test followed by Tukey’s multiple comparison test was performed for parametric data with equal SDs, a Brown-Forsythe and Welch ANOVA followed by Dunnett’s T3 multiple comparisons test was performed for parametric data with unequal SDs, and a Kruskal-Wallis test followed by a Dunn’s multiple comparison test for non-parametric data. When comparing multiple variables across multiple groups, a two-way ANOVA followed by Tukey’s multiple comparisons test was used. Analysis of CFU data was performed on log transformed data. Outliers were determined using the ROUT method (Q=1)^50^.

---

**Table S1.**
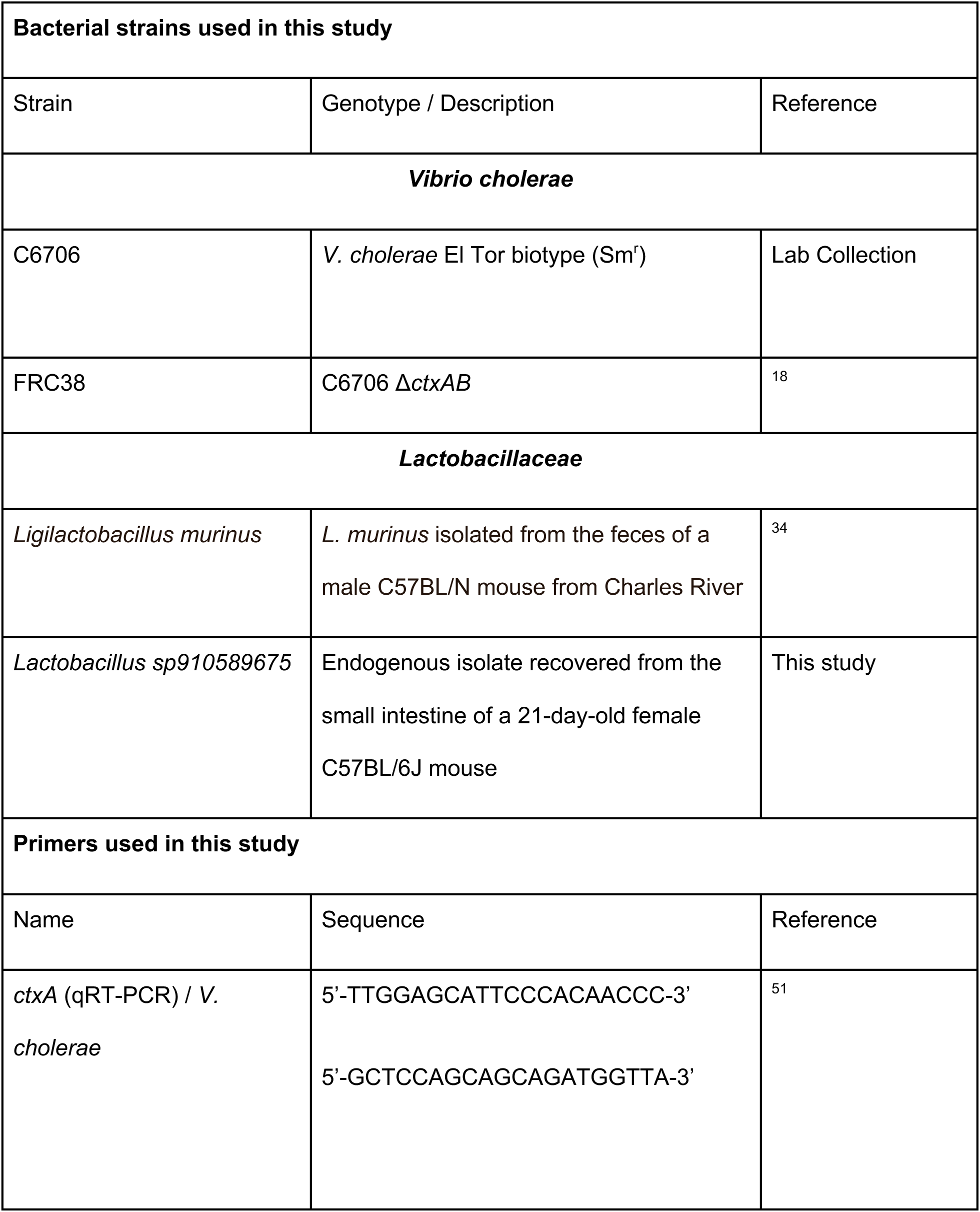
Strains and primers used in this study.

**Figure S1.**
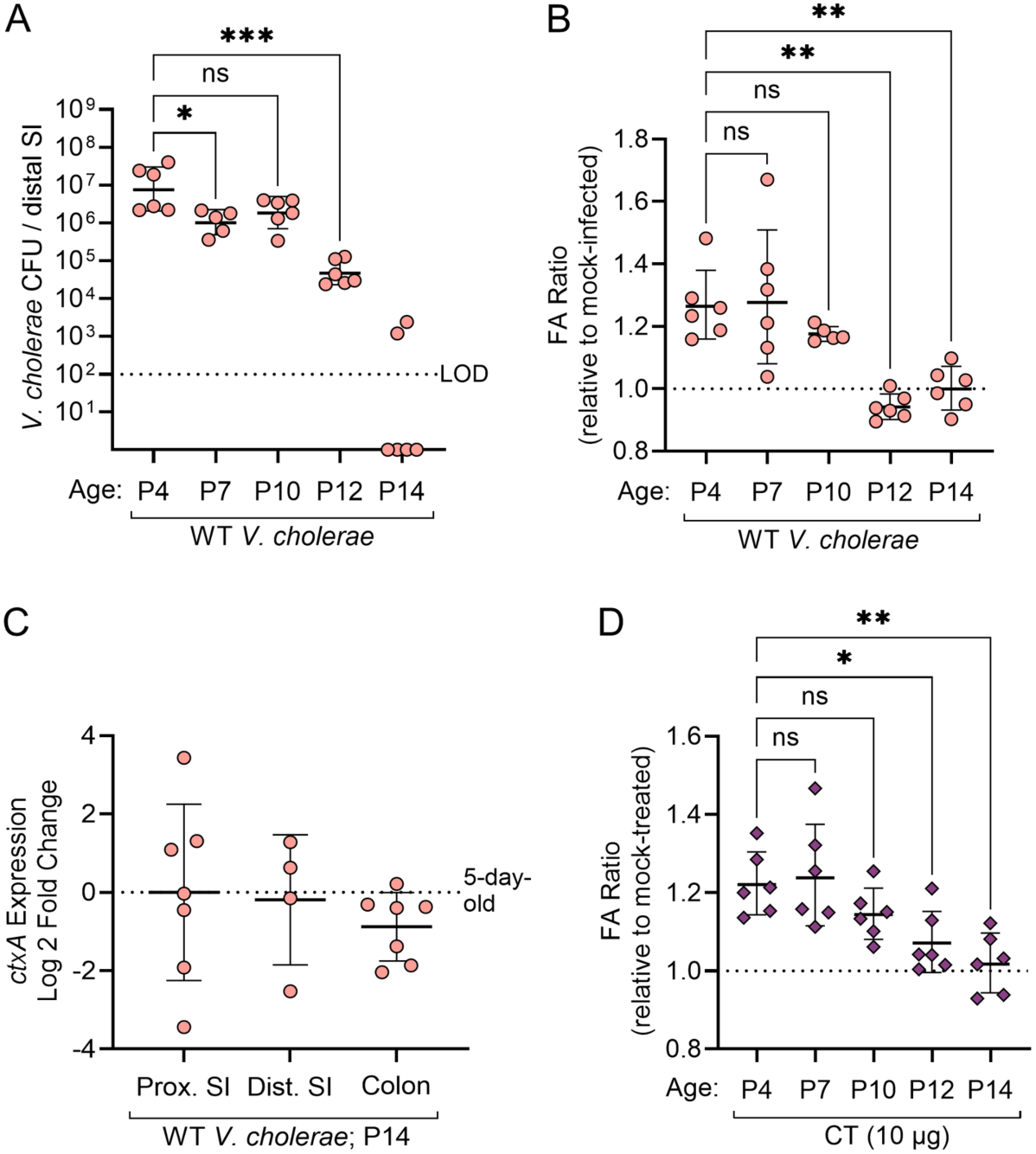
Age-dependent loss of *V. cholerae* colonization and CT responsiveness across mouse backgrounds and intestinal sites. (A–B) CD-1 mice ranging from P4 to P14 were mock-infected with LB broth or infected with 10^7^ CFU WT *V. cholerae*. (A) Distal small intestinal *V. cholerae* CFU. (B) Fluid accumulation ratio relative to mock-infected controls. (C) *ctxA* expression in WT *V. cholerae* recovered from the proximal small intestine, distal small intestine, or colon of P14 C57BL/6J mice, shown relative to expression in P5 mice 22 hours after oral infection. (D) CD-1 mice ranging from P4 to P14 were mock-treated with LB broth or orally treated with 10 μg purified CT. Fluid accumulation ratio is shown relative to mock-treated controls. Each point represents an individual mouse or biological replicate, as appropriate. Bars indicate mean ± SEM. The dotted line in (C) indicates the P5 reference value. Statistical significance in (A–B, D) was determined by Brown-Forsythe and Welch ANOVA with Dunnett’s T3 multiple comparisons test. Statistical

**Figure S2.**
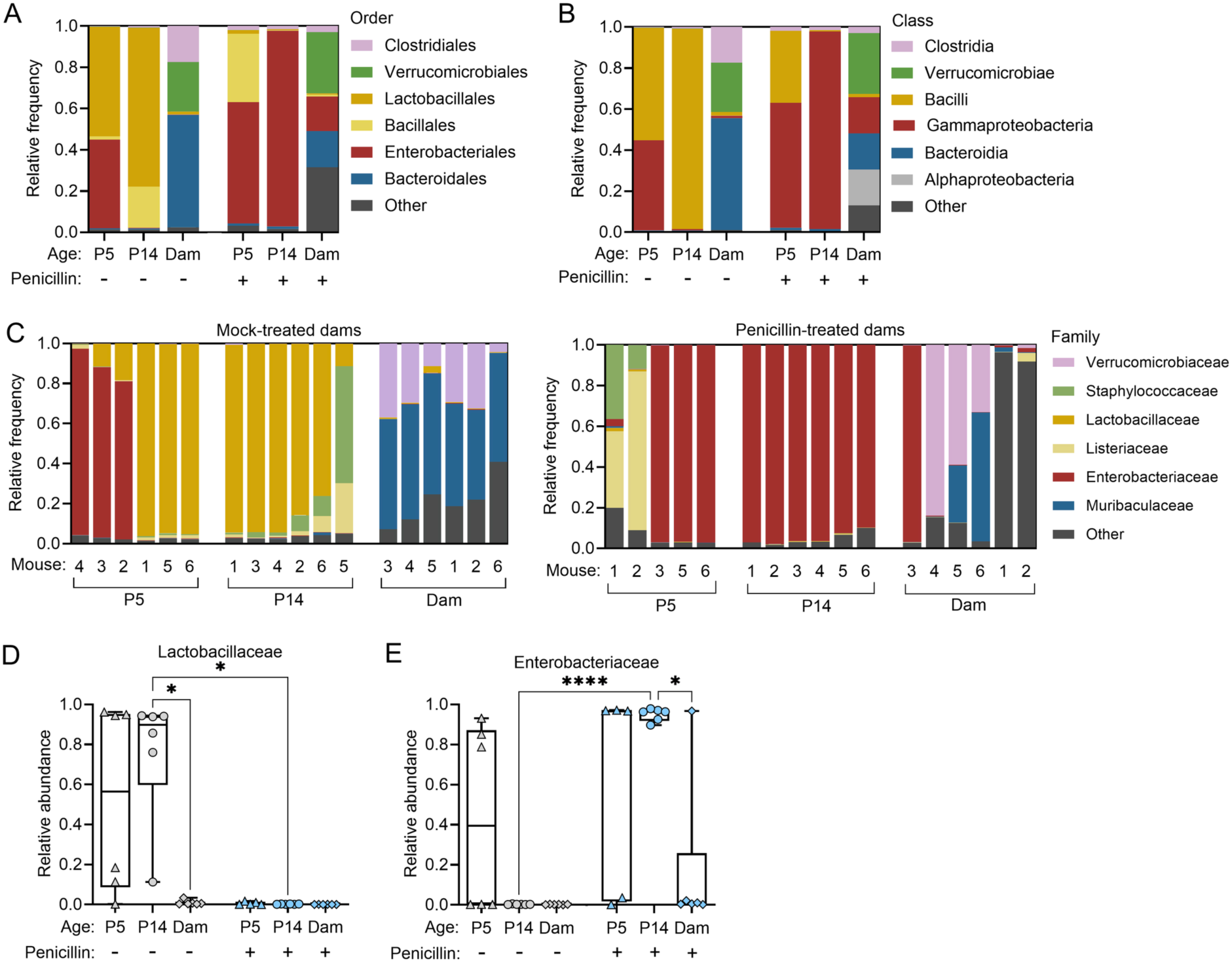
Maternal penicillin alters distal small intestinal microbiota composition in pups and fecal microbiota composition in dams. (A–E) Distal small intestinal bacterial communities from P5 and P14 pups born to untreated or penicillin-treated breeding pairs and fecal communities from corresponding adult dams were analyzed by 16S rRNA gene sequencing. (A) Order-level relative abundance of distal small intestinal or fecal bacterial communities. (B) Class-level relative abundance of distal small intestinal or fecal bacterial communities. Individual-sample family-level relative abundance profiles from untreated (C, left) and penicillin-treated (C, right) groups. Individual-sample family-level relative abundance profiles of Lactobacillaceae (D) and Enterobacteriaceae (E). Each point or bar represents an individual biological sample. Statistical significance in (D–E) was determined by Brown-Forsythe and Welch ANOVA with Dunnett’s T3 multiple comparisons test. ns, not significant; *p < 0.05; **p < 0.01; ***p < 0.001; ****p < 0.0001.

## Notes

### Competing Interest Statement

The authors have declared no competing interest.

